# Preselection of QTL markers enhances accuracy of genomic selection in Norway spruce

**DOI:** 10.1101/2022.11.11.516144

**Authors:** Zhi-Qiang Chen, Adam Klingberg, Henrik R. Hallingbäck, Harry X. Wu

**Affiliations:** Umeå Plant Science Centre, Department Forest Genetics and Plant Physiology, Swedish University of Agricultural Sciences, SE-90183 Umeå, Sweden; Skogforsk, SE-91821 Sävar, Sweden; Skogforsk, Uppsala Science Park, SE-75183 Uppsala; CSIRO National Collection Research Australia, Black Mountain Laboratory, Canberra, ACT 2601, Australia

**Keywords:** Genomic prediction, Marker preselection, GWAS, *Picea abies*

## Abstract

Genomic prediction (GP) or genomic selection is a method to predict the accumulative effect of all quantitative trait loci (QTLs) effects by capturing the linkage disequilibrium between markers and QTLs. Thus, marker preselection is considered a promising method to capture Mendelian segregation effects, especially for an oligogenic trait. Using QTLs detected in the genome-wide association study (GWAS) could improve genomic prediction, including informative marker selection and adding a QTL with the largest effect size as a fixed effect. Here, we performed GWAS and genomic selection studies in a population with 904 clones from 32 full-sib families using a newly developed 50k SNP Norway spruce array. In total, GWAS identified 41 SNPs associated with budburst stage (BB) and the SNP with the largest effect size explained 5.1% of the phenotypic variation (PVE). For the other five traits like growth and wood quality traits, only 2 – 13 SNPs were detected and PVE of the strongest effects ranged from 1.2% to 2.0%. GP with approximately 100 preselected SNPs based on the smallest *p*-values from GWAS showed the largest predictive ability (PA) for the oligogenic trait BB. But for the other polygenic traits, approximate 2000-4000 preselected SNPs, indicated by the smallest Akaike information criterion to offer the best model fit, still resulted in PA being similar to that of GP models using all markers. Analyses on both real-life and simulated data also showed that the inclusion of a large QTL SNP in the model as a fixed effect could improve PA and accuracy of GP provided that the PVE of the QTL was ≥2.5%.

## Introduction

Genomic prediction (GP) or genomic selection using genome-wide dense markers has been widely adopted in animal breeding and extensively studied in crops [1] and tree plant species [2] in the last decade. GP assumes that individual quantitative trait loci (QTLs) are linked with at least one DNA marker. Therefore, linkage disequilibrium (LD) between QTLs and markers plays an important role in genomic prediction efficiency [3]. Most GP results showed that stronger LD between markers and causative mutation resulted in higher accuracy of GPs. It is suggested that training to use multiple generations (tracing LD) or including causative mutations will increase GP efficiency. This is because there is no need to trace the causative mutations with LD markers when the causative mutations are among the genotypes [4]. Therefore, the inclusion of markers tightly associated with a few large-effect QTLs, detected by genome-wide association (GWAS) or validated by gene transformation, could be incorporated into the GP model development [5]. Including such large-effect SNPs as fixed effects in GP modelling is also considered as an ideal approach to increasing GP efficiency, which has been verified by several studies in crops, including simulations [6–8]. GWAS is considered a powerful approach to dissecting the genetic architecture of different traits in animals and plants [9, 10]. Usually, a locus accounting for 1-15% of phenotypic variation is often detected by GWAS if population sizes are large enough, from a few hundred to a few thousand in plants [11], and a few thousand to a few million in humans [12].

Several studies in trees have shown that a few thousand randomly selected SNPs may capture most of the variation, and similar GP efficiency as using all available markers [13–16]. It has also been reported that marker preselection could slightly to moderately improve GP accuracy in tree species [17–19]. In theory, GP would like to capture all QTL effects by LD between markers and causative loci. Providing that one or two of several strongly linked markers for each QTL is selected, the model should almost capture the QTL effects. Meanwhile, if one can select the tightly linked markers or mutants themselves through GWAS for GP, it is possible to reduce the cost of genotyping while maintaining or improving the accuracy of GP. Recently, following the development of high throughput genotyping techniques and tools, more than several hundred thousand markers have been commonly and easily produced by several genotyping platforms, such as SNP array, exome capture, and genotyping-by-sequencing (GBS). Thus, marker preselection could become a very useful and common pre-step for GP.

Norway spruce (*Picea abies* (L.) Karst.) is one of the most important economic species in Europe, especially in the Nordic countries and Northwestern Russia [20]. The current breeding program of Norway spruce, similar to other conifer species, mainly focuses on the use of additive genetic effects (i.e. breeding values). However, the non-additive genetic effects are considered important if a clonal deployment is considered as a deployment strategy in the future [21–23]. Recent several studies on the cost, benefit and genetic diversity with the deployment of clonal forestry for Norway spruce indicated that a considerable productivity increase of planted forests could be achieved with an acceptable genetic diversity [23–30].

In Norway spruce breeding, traditional breeding values, dominance, and epistatic genetic effects were predicted for seedling or clonally propagated progenies based on the theoretical expectation using pedigree data [23]. When genome-wide dense markers are available, the estimates of genetic parameters for additive, dominance, and epistatic effects may be more accurate since the real genomic relationships were captured by alleles [31, 32]. Several studies in tree species have demonstrated an increase in the accuracy of genetic parameters estimates using genomic models [33–35]. The accurate estimation of non-additive effects, as well as additive effects, should therefore be an important objective to increase genetic gains and improve the efficiency of the Norway spruce tree breeding program.

In this study, we explored the use of detected GWAS QTLs by including the most closely associated SNPs in GP for tree breeding value prediction through empirical experiments and simulations. The detailed aims of this study were to 1) dissect the total genetic component into additive and non-additive variances using genomic-based relationship matrix models and compare with the corresponding dissection using traditional pedigree-based relationship models; 2) test the efficiency using different number of SNPs, different number of clones per family with informative maker preselection; 3) optimize the training population size and population structure for the current Swedish breeding population; and 4) test whether the GP could be improved by including the most significant GWAS marker as a fixed effect in the GP model.

## Results

### Genetic parameter comparisons between models

We compared the goodness of fit of four models (equations (1–4), Table 1 and Table S1). For all traits, the GBLUP-AR model had the smallest Akaike information criterion (AIC) value, except for frost damage (FD) with a zero non-additive variance (i.e. GBLUP-A). This indicates that the fitting of GBLUP-AR was generally better than all other models, both pedigree-based and genomic-based, and implies that the genetic parameters of the GBPLUP-AR models should be better estimated than for the other models. Based on the AIC, we did not see that models with first-order epistatic effect terms (PBLUP-ADR-xx and GBLUP-ADR-xx) showed any better fit than the best pedigree- or genome-based models in that respect (usually PBLUP-AR and GBLUP-AR, respectively).

**Table 1.**
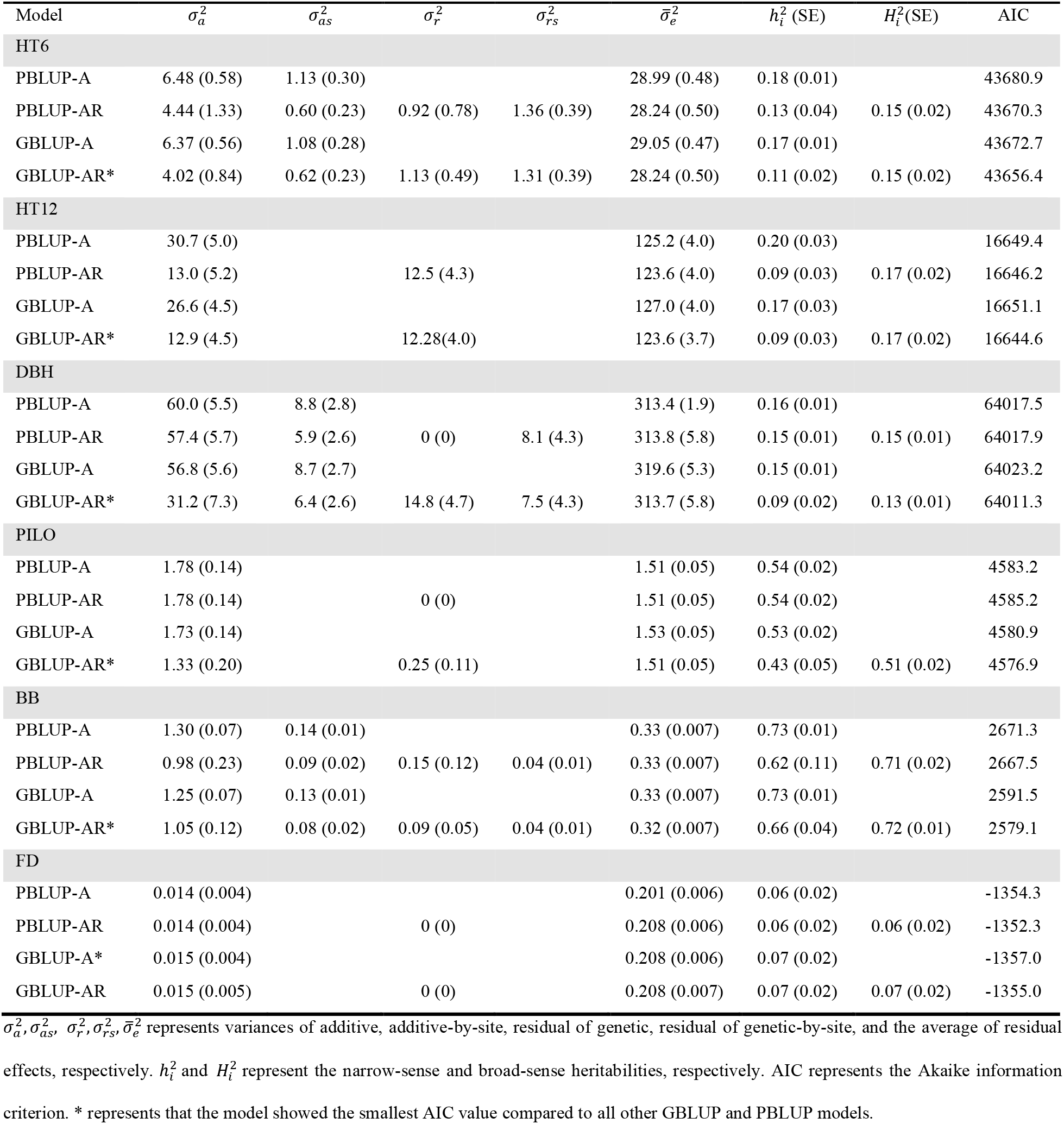
Ratios of additive, dominance, and residual of genetic variance component to the total phenotypic variance for different PBLUP and GBLUP models.

### Estimates of variance component and heritability

We found that the additive genetic variance under PBLUP-AR and GBLUP-AR decreased compared to that under PBLUP-A and GBLUP-A for HT6, HT12, DBH, and BB, respectively (Table 1). Given that the PBLUP-AR and GBLUP-AR exhibited the best fit for all these traits, the previously observed patterns indicate that the additive genetic variance in the PBLUP-A and GBLUP-A may be inflated due to the inadvertent inclusion of non-additive effects within 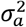. For the growth traits (HT6, HT12 and DBH), the estimates of the non-specific non-additive genetic variance 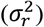 were substantial in size relative to the additive genetic variance estimates 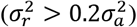 under the model with lowest AIC (GBLUP-AR). For BB and PILO the non-additive variance estimates were relatively small in comparison to the additive variance 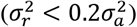. Finally, FD did not exhibit any non-additive effects regardless of whether the pedigree- or genome-based matrix models were used. Taken together, we may conclude that non-additive effects are important for all growth traits, whereas non-additive effects had a limited impact on non-growth traits BB and PILO and were absent in FD. Estimates of dominance variance were generally small and non-significant and most of the first-order epistatic interactions under PBLUP-ADR-xx and GBLUP-ADR-xx were in boundary close to zero based on Equation [4] (Table S1). Thus, we do not discuss the result of the PBLUP-ADR, GBLUP-ADR, PBLUP-ADR-xx and GBLUP-ADR-xx models any further.

### Summary of Norway spruce 50K SNPs array, LD decay, and association mapping

For the *Piab50K* SNP array, 41,236 of the total 47,445 SNPs were mapped onto each of the 12 chromosomes in the Norway spruce genome v2 (In preparation). The number of SNPs in each chromosome varied from 3158 to 3991 SNPs (Table S2). The physical extent of LD (*r*^2^ > 0.2) within each chromosome varied from 33.7 kb in chromosome 2 to 54.6 kb in chromosome 9, with an average of 42.9 kb for the whole genome based on the SNP array in the studied full-sib family population (Fig. S1). The family clusters were clearly separated by the first two principal components (Fig. S2). In total, GWAS identified 41, 11, 2, 4, 4, 11, and 13 SNPs as having a significant effect on BB, DBH, FD, HT12, HT6, and PILO, respectively, under a false discovery rate (FDR) of 0.05 (Fig. 1b). The largest effect sizes for trait-associated SNPs explained 5.1, 1.6, 1.7, 2.0, 1.2, and 1.4% of the phenotypic variation of clonal means in BB, DBH, FD, HT12, HT6, and PILO, respectively (Table S3). There were six highly significant SNP-trait associations with considerable PVE (>2.5%) for BB, indicating oligogenic nature whereas such observations were not made for the other traits.

**Fig.1.**
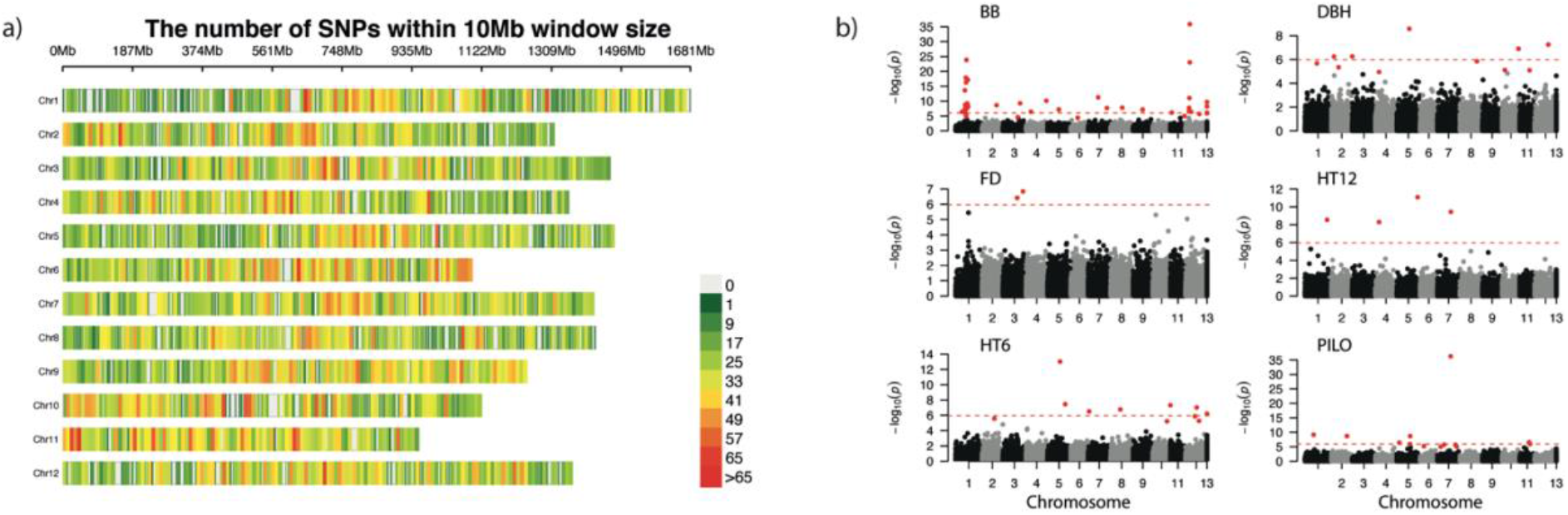
Marker density of this SNP array and Manhattan plots. a) Marker density of the 50k Norway spruce array based on a 10Mb window size for each of 12 chromosomes. b) Manhattan plots for six traits. The red dashed line represents the significant threshold of *p* = 1.7 × 10^−6^ after the Bonferroni correction. The red dots represent that the SNPs passed the false discovery rate test threshold of 0.05 based on Benjamini and Hochberg (1995). SNPs which were not mapped into the Norway spruce genome v2 (In preparation) were grouped into a region assumed as chromosome 13 in this study. BB, bud burst stage; DBH, diameter at breast height; FD, frost damage; HT12, tree height at field tree age 12; HT6, tree height at field age six; PILO, Pilodyn penetration.

### Predictive ability comparisons between PBLUP and GBLUP

We initially compared predictive abilities (PAs) estimated from cross-validation under PBLUP and GBLUP models with the square root of the clone mean narrow-sense heritability (*h_c_*, Table 2 and Table S4) because it can be interpreted as the theoretical PA and accuracy for mass selection based on phenotypic clonal means. In agreement with previous comparisons between PBLUP and GBLUP in terms of AIC, we observed that PA-estimates of GBLUP (0.23 – 0.67) were consistently higher than corresponding PA-estimates for PBLUP (0.20 – 0.61). The increases in PA under GBLUP were the greatest for HT6 and BB. For example, PA of GBLUP for HT6 increased by 15.8% when compared to PBLUP. However, the PA-estimates for GBLUP in a 10-fold cross-validation scheme (phenotypic data absent) were in turn consistently and considerably lower in comparison to the corresponding theoretical PAs in the presence of phenotypic data (*h_c_* in the range 0.37 – 0.89).

**Table 2.** Overall predictive ability (PA) for different types of pedigree- and genome-based prediction models following a 10-fold cross-validation procedure.

| Model | HT6 | HT12 | DBH | PILO | BB | FD |
| --- | --- | --- | --- | --- | --- | --- |
| $h_c$ | 0.72 | 0.42 | 0.69 | 0.87 | 0.89 | 0.37 |
| PBLUP-C | 0.38 (0.01) | 0.22 (0.01) | 0.44 (0.01) | 0.62 (0.01) | 0.61 (0.01) | 0.21 (0.01) |
| GBLUP | **0.42 (0.01)** | 0.24 (0.01) | **0.44 (0.01)** | **0.63 (0.01)** | 0.67 (0.01) | 0.23 (0.01) |
| GBLUP-S | 0.35 (0.01) | 0.22 (0.01) | 0.37 (0.01) | 0.58 (0.01) | **0.72 (0.01)** | 0.20 (0.01) |
| GBLUP-F | **0.42 (0.01)** | **0.26 (0.01)** | **0.44 (0.01)** | **0.63 (0.01)** | 0.70 (0.00) | **0.25 (0.01)** |
| Accuracy | 0.58 | 0.62 | 0.64 | 0.72 | 0.81 | 0.68 |

The prediction models showing the highest PA for a trait have their PA-estimated highlighted in *italic bold*. The $h_c$ is the square root of the clonal mean narrow sense heritability ( $h_c^2$ ) based on PBLUP-AR model. PBLUP-C is the traditional pedigree-based best linear unbiased prediction (BLUP) including marker-based pedigree correction; GBLUP is the genomic-based BLUP; GBLUP-S is genomic-based BLUP and with marker-preselection based on the smallest $p$ -values from genome-wide association analyses. In this table, the 100 smallest $p$ -value SNPs were preselected for the budburst stage (BB) and 2000 SNPs were preselected for the other traits. GBLUP-F is a genomic-based BLUP model plus the SNP with the smallest $p$ -value fitted as an additional fixed regression effect. The accuracy for the model showing the highest PA is here roughly estimated as $PA/h_c$ .

### The impact of the number of SNPs included in the G-matrix

For all traits, the PA of the GBLUP model increased from 25 randomly selected SNPs to ca. 4000 SNPs (Fig. 2a) and plateaued when even more SNPs were selected. Similarly, the standardized AIC values of the GP model decreased from the model with 25 SNPs to ca. 4000 SNPs and then the AIC stabilized after that (Fig. 2b). For example, for BB, PA under the GBLUP-model increased from 0.36 using 25 SNPs to 0.63 using 4000 SNPs in the estimation of a genome-based relationship matrix.

**Fig. 2.**
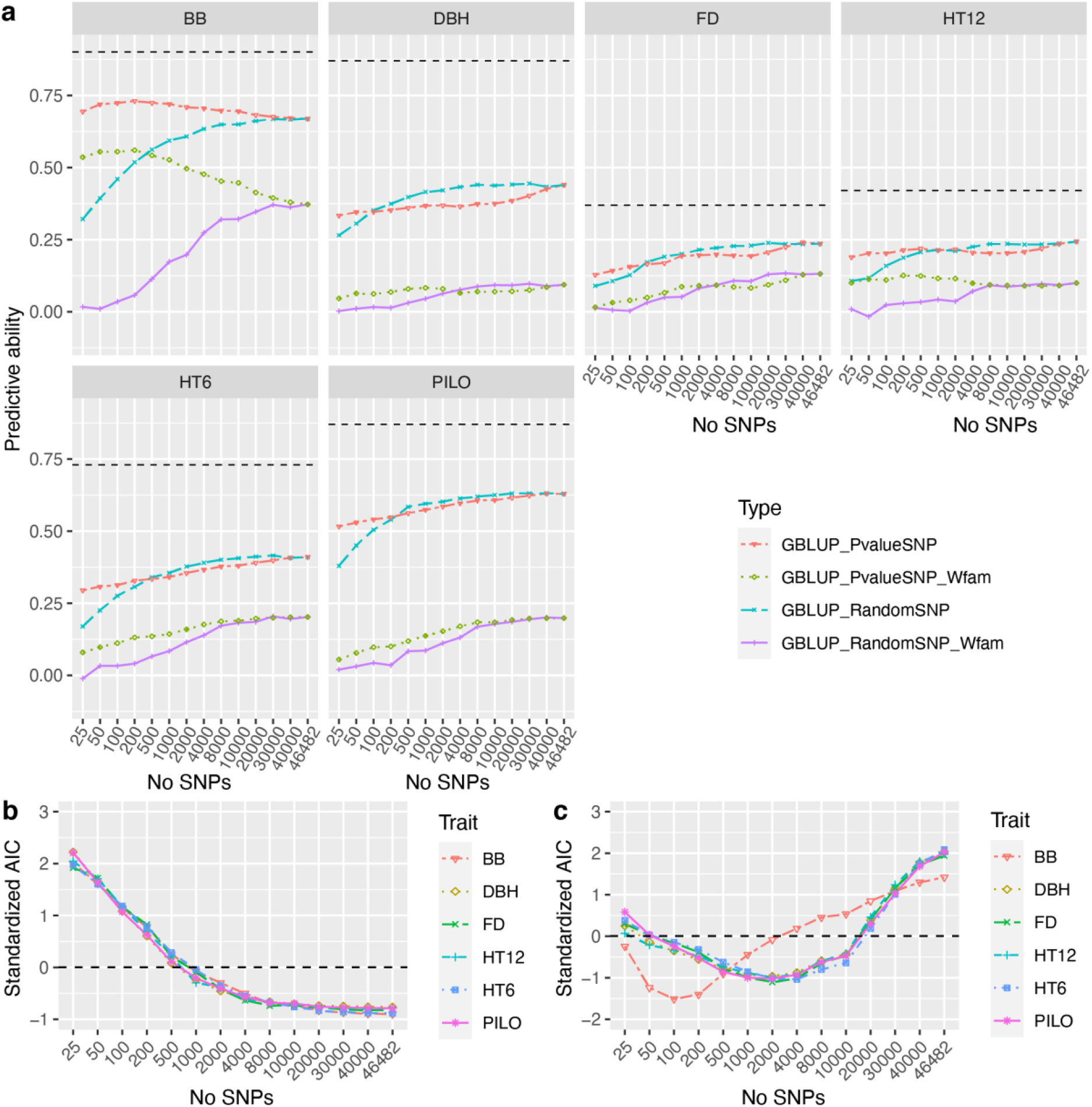
Predictive ability estimates (PA) for six traits using GBLUP-A models based on 10 repeats of 10-fold cross-validations (n=100) and 904 clone means. a) PAs for trait’s phenotypic values based on GBLUP-A using different numbers of SNPs randomly selected (blue longdashed line), SNPs selected based on lowest *p*-values in GWAS performed on the training dataset using BLINK method (red twodashed line), and PAs of within family variation (clone mean - family mean) using different number of SNPs based on both randomly selected (purple solid line) and the lowest *p*-values based on GWAS (green dotted line). The black horizontal dashed line in a) is the square root of narrow-sense clone mean heritability estimated based on the PBLUP-AR model. Matching to GBLUP-A models in a), the standardized Akaike information criterion (AIC) values for each model is shown in b) based on randomly selected SNPs and in c) based on SNPs selected by smallest *p*-values.

When marker preselection was based on GWAS-analyses, the resulting prediction models showed two types of trends for the traits. First, for BB, the genomic prediction model (GBLUP-S) with 100-200 preselected SNPs with the smallest *p*-values obtained a higher PA than that using all other numbers of SNPs (Fig. 2a). To check if GP with a marker preselection by GWAS captured more Mendelian segregation effects, we also calculated the correlation between estimated breeding value deviations from family mean of EBVs and within-family phenotypic variation (clone mean-family mean). We found that GBLUP-S using 100-200 informative markers captured more of the phenotypic variation of BB and exhibited the lowest AIC value (Fig. 2c). This suggests that using all available SNPs for a trait indicated to be oligogenic may introduce misleading noise in the prediction models.

Second, for the other five traits (DBH, FD, HT12, HT6, and PILO), we found that the PA of GP using a preselected set of SNPs with the smallest *p*-values did not outperform random marker selection unless the numbers of SNPs selected were less than a few hundred. Furthermore, for these traits preselecting more markers beyond a few hundred still increased the PA with the highest value being reached when all markers were used (Fig. 2a). For traits DBH, FD, HT12, HT6, and PILO, within-family PA for marker selection based on low *p*-values was higher than for random selection when smaller numbers of markers were selected all ranging from 25 SNPs to 2000 SNPs. But random selections of markers showed ca. equal and higher within-family PA than GWAS-based preselection for these traits when more than 10000 SNPs were selected. The trends of PA when selecting more and more markers for DBH, FD, HT12, HT6, and PILO were not obvious where a flat within-family PA optimum for GWAS-based preselection was observed for HT12 and DBH (at 1000 and 200 markers respectively) whereas for the other traits more markers resulted in higher within-family PA. AIC values for all traits except BB showed a similar trend (Fig. 2c) where the lowest AIC value occurred at 2000-4000 preselected markers.

### Predictive ability comparisons between advanced GBLUP models

As previously indicated in Fig. 2, only BB showed appreciable increases in overall PA from 0.61 for GBLUP to 0.72 for GBLUP-S when only 100 high-significance SNPs in GWAS were included in the *G_a_*-matrix (Table 2). For all other traits the GBLUP-S model including 2000 high-significance SNPs showed similarly or lower PA than the conventional GBLUP including all SNPs. In addition, we attempted the fitting of a single SNP showing the smallest *p*-value (the highest significance) in GWAS as an additional fixed regression effect in the model (GBLUP-F). For HT12, BB, and FD, the GBLUP-F model produced a slightly higher PA than GBLUP (by 0.02 – 0.03) but for DBH, HT6, and PILO, GBLUP-F and GBLUP showed similar values. Furthermore, we calculated an approximation of cross-validation accuracy in the absence of phenotypic data as the PA of PBLUP and GBLUP divided by *h_c_* and we found that the overall accuracy of GP for all traits ranged from 0.58 for HT6 to 0.81 for BB.

### Simulations of including a large effect SNP as fixed in the GBLUP

To verify whether including a major gene locus as a fixed effect in the GBLUP model (GBLUP-F) would increase PA of GP, we also performed 10 repeats of 10-fold cross-validations on simulated data where single SNPs exerting a range of PVE were included (Table 3). Simulation data is easier to interpret than real-life data since it offers the possibility to robustly estimate GP accuracies merely by calculating the correlations between predicted and true breeding values. Results showed that the overall accuracies of the GBLUP-F model were higher than that of GBLUP when provided a major-effect SNP with PVE>=2.5% (0.66-0.71 and 0.60-0.63 for GBLUP-F and GBLUP respectively). Following a similar trend, the overall PA of the GBLUP-F model increased from 0.31 to 0.35 for a fixed SNP explaining 0% to 5% of the phenotypic variation whereas no increasing trend was observed for the model without the major QTN included.

**Table 3.** Predictive abilities and accuracy estimates with genomic prediction model with and without adding an SNP with the largest effect size as a fixed effect (GBLUP and GBLUP-F) based on simulated data using a trait with a heritability of 0.25 and evaluated using 10-fold cross-validations.

| PVE (%) | Overall |  | Within-family |  | Overall accuracy |  | Within-family accuracy |  |
| --- | --- | --- | --- | --- | --- | --- | --- | --- |
|  | predictive ability |  | predictive ability |  |  |  |  |  |
|  | - QTN | + QTN | - QTN | + QTN | - QTN | + QTN | - QTN | + QTN |
| 5 | 0.31 (0.06) | 0.35 (0.06) | 0.03 (0.07) | 0.17 (0.07) | 0.60 (0.05) | 0.71 (0.04) | 0.11 (0.08) | 0.42 (0.06) |
| 2.5 | 0.33 (0.07) | 0.34 (0.06) | 0.07 (0.08) | 0.11 (0.07) | 0.63 (0.06) | 0.66 (0.05) | 0.12 (0.09) | 0.29 (0.07) |
| 1.25 | 0.31 (0.06) | 0.31 (0.07) | 0.02 (0.06) | 0.07 (0.07) | 0.63 (0.05) | 0.64 (0.05) | 0.09 (0.07) | 0.20 (0.08) |
| 0.25 | 0.30 (0.07) | 0.30 (0.06) | 0.05 (0.06) | 0.06 (0.07) | 0.61 (0.05) | 0.60 (0.06) | 0.11 (0.07) | 0.13(0.08) |
| 0 | 0.32 (0.06) | 0.31 (0.07) | 0.05 (0.07) | 0.05 (0.07) | 0.62 (0.06) | 0.61 (0.07) | 0.11 (0.08) | 0.10 (0.08) |

PVE is the percentage of phenotypic variance explained by a large effect size SNP. QTN represents the quantitative trait nucleotide with a large effect size where - and + signifies the absence (GBLUP) and presence (GBLUP-F) respectively, of this QTN in the genomic prediction model as a fixed effect. The value in parenthesis indicates the standard deviation across replicate simulations.

The within-family accuracy of the GBLUP-F model increased from a very low level (0.10) when the fitted fixed-effect SNP explained 0% of the variation (i.e. fitting a false-positive QTL) to 0.42 for a SNP with a PVE at 5% (true major gene, Table 3). In contrast, for the model where the assumed major SNP was not fitted as a fixed effect, within-family accuracies remained at very low levels (0.09-0.11) regardless of the PVE of the major SNP. The trends of within-family PA were similar as those for accuracy but were all lower due to the environmental noise that always influences PA-estimates. The above results indicate that including a large-effect SNP (PVE>=2.5%) as a fixed effect in the GBLUP model may improve the accuracy and PA, especially with respect to predictions within family.

### Number of clones per family

To test the effect of the number of clones per family available to the training dataset on the PA using PBLUP and GBLUP (Fig. 3), we sampled 5, 10, 15, 20, 25, or 30 clones from each of the ten largest families as a training data set and the rest of clones in those families as a validation set. We found that the PA of both PBLUP and GBLUP consistently increased from 5 clones per family to 30 clones per family for all traits except for PBLUP for BB where an optimum was reached at 20 offspring clones per family. Based on the trends and with GP in mind (GBLUP), more than 30 clones would be better regardless of the trait under study. Following the increase of the number of clones per family, GBLUP showed a higher PA than PBLUP, except for DBH where PA-estimates were similar.

**Fig. 3.**
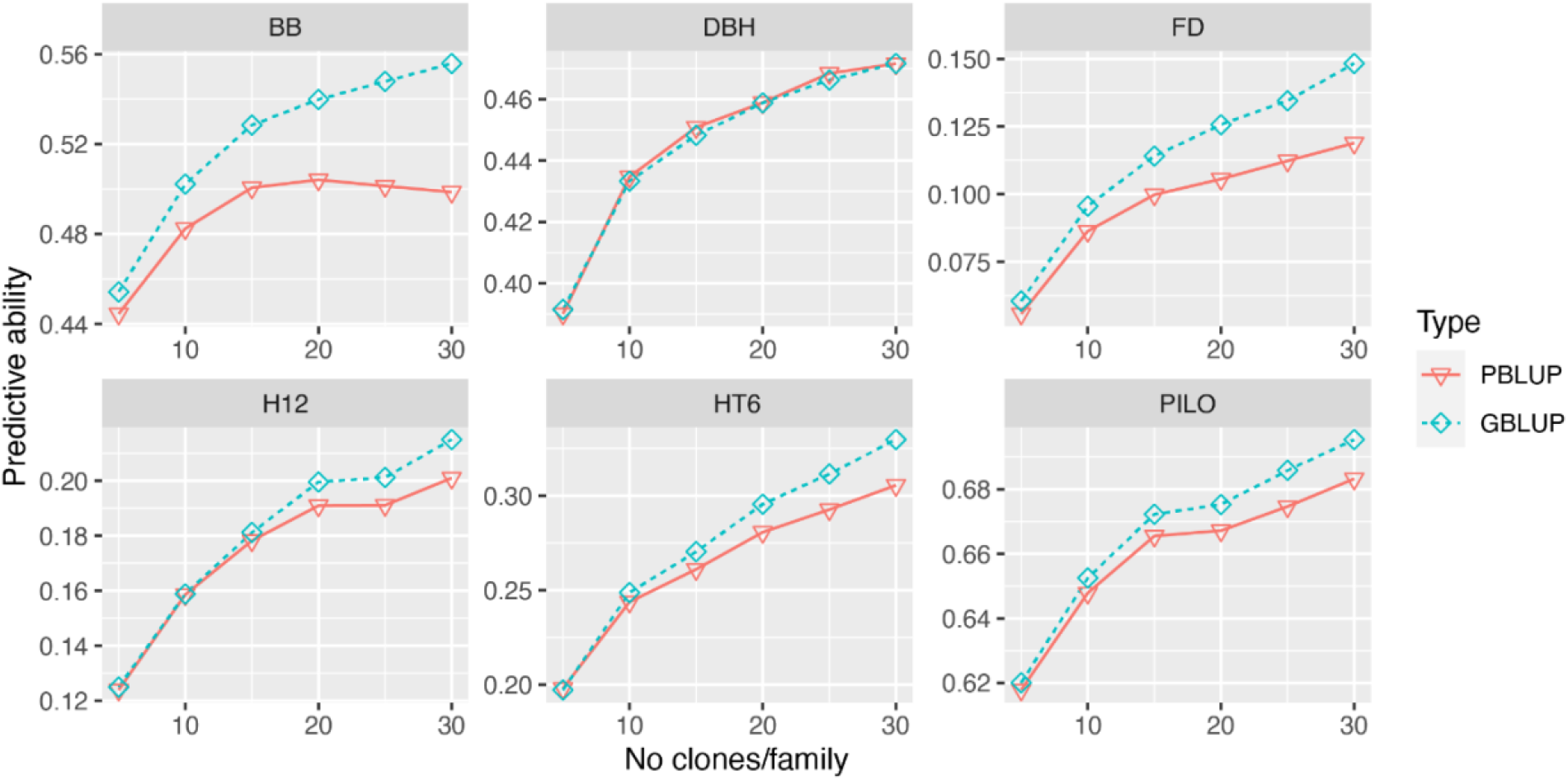
Predictive abilities (PAs) for six traits using PBLUP-A and GBLUP-A models using all available SNPs but employing different number (5, 10, 15, 20, 25, and 30) of clones per family for model training leaving the remainder number of clones for validation purposes. Only ten families comprising the largest numbers of offspring were included in this analysis.

### Number of clones per family and marker preselection by GWAS also affect the within-family variation

To further test whether combining GWAS *p*-value based marker preselection and the number of clones per family would affect the predictive ability (PA) within families, we performed 10-fold cross-validations using GBLUP within the largest ten families only (Fig. 4). We found that the PA within the ten largest families using GBLUP with GWAS-based preselection of a number of markers produced similar trends (Fig. 4a) as the corresponding PA values estimated for the population as a whole (Fig. 2a). In similarity to the results of the whole population, GBLUP of BB with 100-200 preselected SNPs produced higher predictive ability and lower AIC than using a lesser or greater number of SNPs for *G_a_*-matrix calculation (Fig. 4a and b). For the other traits, 2000-4000 preselected SNPs produced the lowest AIC in agreement with the whole-population analysis. The trends for within-family PA, although less obvious, indicated that some sort of SNP preselection resulted in higher PA than uncritically using all SNPs in the model.

**Fig. 4.**
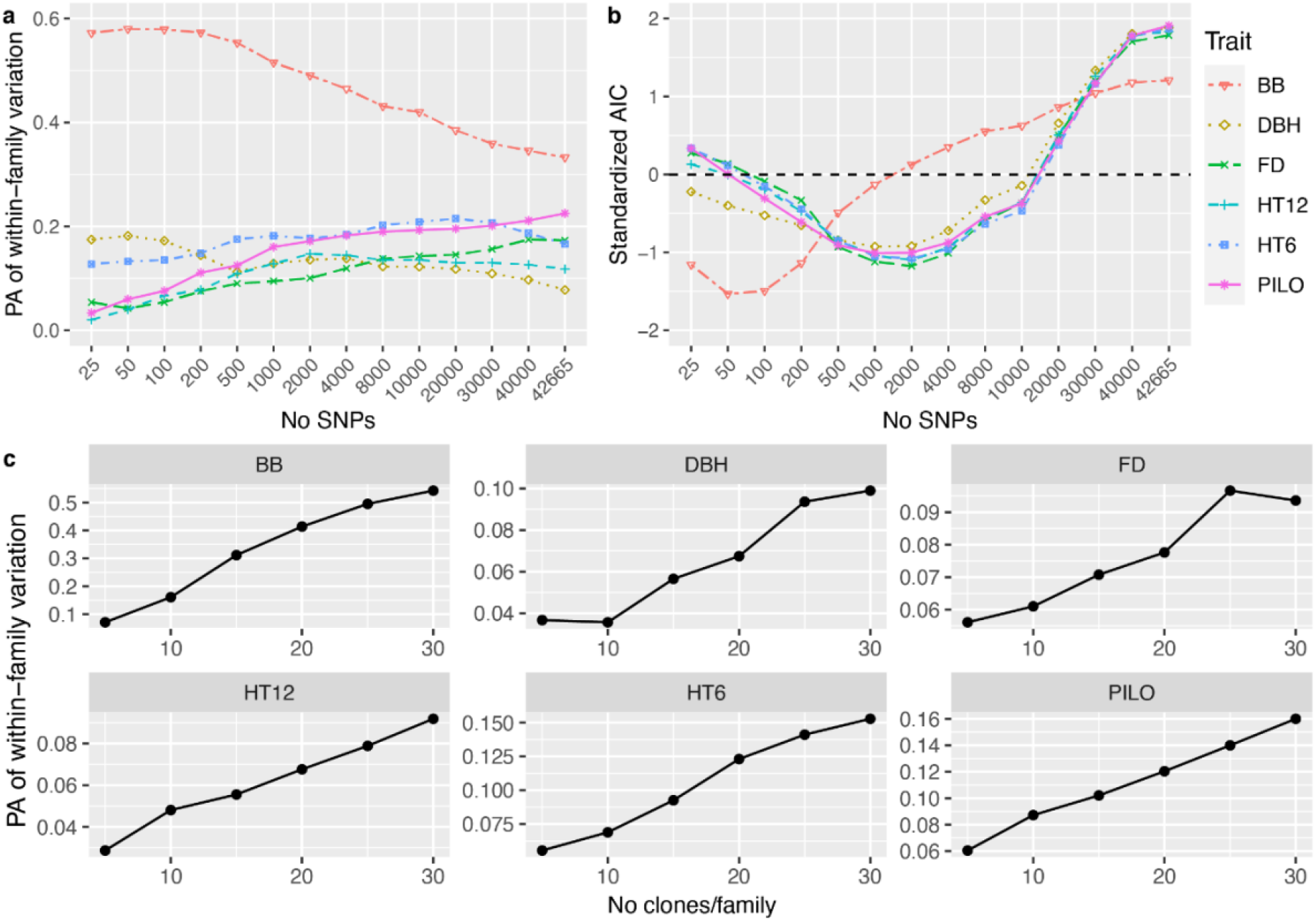
a) Predictive ability (PA) for within-family variation using 10-fold cross-validation for the ten largest families with 482 clones. b) the standardized Akaike information criterion (AIC) values for different simulation strategies based on different numbers of SNPs with the smallest p values from GWAS. c) PA for within-family variation based on the GBLUP model with G matrix estimated from 100 SNPs for budburst and 2000 SNPs for the rest of the traits based on cross-validation GWAS in the ten families.

These results could imply that more within-family variation linked to Mendelian segregation could be captured by preselecting influential markers for GP and that the capture of such segregation variation is likely easier in a population where families are fewer and larger thus offering a higher average relationship. Based on the marker preselection results for within-family prediction (Fig. 4c), we also found that PA increased quickly from 5 clones per family to 30 clones per family for all traits, except for FD, which showed a slight decrease for within family prediction after 25 clones per family. For example, PA within-family variation for BB increased from 0.07 at family size 5 to 0.54 when 30 clones per family were available for model training.

In practical breeding, GP may be performed for a new full-sib family which does not offer close relationships with existing families in a training dataset. Thus, we also evaluated one special type of cross-validation in which the clonal mean data for each of 32 families were in turn held out from the GBLUP training dataset, while the family for which clonal mean data was missing was used for the prediction validation (across-family validation). As previously, we performed the GBLUP with 1) all markers for *G_a_*-matrix calculation (GBLUP), 2) 100 preselected SNPs for BB and 2000 preselected SNPs for the rest of the six traits based on significant SNP-trait associations (GBLUP-S), and 3) all markers for the *G_a_*-matrix plus the marker with the highest GWAS significance fitted as a fixed regression effect in the model (GBLUP-F). We found that marker preselection increased PAs for some traits (HT12, PILO, BB, and FD), but not for others (HT6 and DBH). The PA-estimates across families were very low for most traits (0.03 to 0.11) but for BB the GBLUP PA was 0.19, increased to 0.24 when fitting the most powerful SNP as a fixed effect (GBLUP-F), and increased further to 0.25 when using GBLUP-S (Table 4).

**Table 4.** Predictive ability for clone mean phenotypic variation and their standard errors in parenthesis for different models where the validation procedure was performed across different families.

| Model | HT6 | HT12 | DBH | PILO | BB | FD |
| --- | --- | --- | --- | --- | --- | --- |
| GBLUP-All | 0.11 (0.05) | 0.03 (0.04) | 0.05 (0.04) | 0.05 (0.04) | 0.19 (0.05) | 0.03 (0.04) |
| GBLUP-S | 0.03 (0.04) | 0.07 (0.04) | -0.02 (0.04) | 0.08 (0.03) | 0.25 (0.06) | 0.04 (0.03) |
| GBLUP-F | 0.09 (0.04) | 0.02 (0.04) | 0.04 (0.04) | 0.04 (0.04) | 0.24 (0.06) | -0.01 (0.04)) |

In this scenario (GBLUP), each of the 32 full-sib families was a validated set and a training set using the rest of the 31 families. In total, 32 cross-validations were repeated to calculate the predictive ability for phenotypic variation. GBLUP-All, GBLUP model with all SNPs; GBLUP-S, GBLUP model with preselected SNPs (100 SNPs for budburst stage (BB), 2000 SNPs for the rest of the traits); GBLUP-F, GBLUP model with the most significant *p*-value SNP as a fixed effect.

## Discussion

### Genomic-based BLUP model could enhance the accuracy of the estimated additive and non-additive genetic variances

In tree or conifer breeding programs, a control-pollinated clonal test provides an opportunity to simultaneously estimate the additive, non-additive, dominance, and epistatic variances using pedigree. However, most heritability estimates in many traditional tree breeding programs were based on open-pollinated or control-pollinated progeny trials without vegetative propagation. In this context, such estimates of additive genetic variation may be biased due to the difficulty of separating the additive genetic variance from parts of the dominance and epistatic variances [36]. Using vegetatively propagated material in combination with a genome-based model, the additive and dominance variances may still show bias if the model does not include a residual genetic effect [32, 35]. Potentially, genome-based models could capture and discriminate between the different sources of the non-additive genetic variance such as dominant effects within a locus and epistatic interaction effects among loci. Among different GP models, GBLUP-AR showed the smallest AIC values for most of the traits, indicating that non-additive genetic effects are significant for all traits, with frost damage being the only exception. Also, the fact that GBLUP-AR model AIC values for these traits were systematically lower than the corresponding PBLUP-AR AIC, implies that GP models perform better with respect to separating additive and non-additive variances from each other than pedigree-based models. For example, PILO showed non-significant and negligible non-additive effects under PBLUP-AR but showed nonetheless significant non-additive effects under the GBLUP-AR model, indicating that the GBLUP-AR model with a realized relationship matrix could better capture and separate the additive genetic variation thus improving genetic parameter estimates.

### Significant marker-trait associations and genomic prediction

Recently, several studies on tree species reported that selecting markers with particular influence over a trait could improve PA [16]. Tan and Ingvarsson (2022) [19] reported that a careful 1% preselection of markers could improve the estimate of heritability and GP in a *Eucalyptus* population. For *Pinus contorta* Douglas ex Loudon var. *latifolia*, Cappa et al. (2022) [17] reported that selecting informative markers, in particular markers capturing ancestry/population structure, can improve PA.

The recently developed 50k SNP array for Norway spruce included several QTLs per trait detected in our previous GWAS [37–40]. In this study, GWAS identified 44 associated SNPs for the budburst stage and one SNP (MA_12842_2274, Table S3), having the second highest PVE (>4%) of all SNP-BB associations, was located within 400 base pairs from a QTL for BB previously detected by GWAS in a different population of ca. 4000 individuals [40]. We also observed two to 13 SNPs significantly associated to the other five traits, offering the opportunity to test if including the most significant SNP in GP model improves PA.

### Marker preselection and inclusion of a large effect QTL as a fixed effect could enhance GP predictive ability

For BB, we found that a *G_a_*-matrix built from 100 preselected markers resulted in GBLUP-models having lower AIC values and being better at predicting the genetic value in the absence of phenotypic data than did a *G_a_*-matrix model using all available markers. Such preselected SNPs were also observed to capture a considerable amount of within-family variation/Mendelian segregation effects. However, for other more polygenic traits, genomic models using ca. 4000 preselected markers showed a comparable PA compared to the model using all markers, even though such models exhibited lower AIC values. This indicates that the preselection of influential markers is more likely to be successful when applied to a limited set of markers showing highly significant associations to an oligogenic trait where a relatively limited number of QTLs are likely to exhibit considerable individual effects (PVE > 1%).

In this study, we also investigated a realized GP model for the BB where a large-size QTL with a PVE of ca. 5% was explicitly included as a fixed regression effect and we observed a 4.4% improvement in overall PA (Table 2) in comparison to a GP model without such a modification. Such enhancement was also observed in several empirical crop studies [41, 42] and a simulation study conducted by Bernardo (2014) [6]. Our finite-locus simulations also showed a similar result with 13% and 18% of improvement, in terms of PA and accuracy respectively, for the model including a locus of large effect size (PVE at 5%) as a fixed effect compared with the model without this model term. The model improvement was particularly notable when the objective was the capture of within-family Mendelian segregation variation (Table 3). However, if the QTL PVE was less than ca. 1.25%, the simulated data analyses did not indicate any such advantage for the model including the locus as a fixed effect. Based on more than dozens of GWAS results in tree species [11], SNPs detected with a PVE >1.25% are not uncommon, indicating that GP appropriately utilizing a large-size QTL should be very useful to improve PA and accuracy. This may indicate that GP using GWAS results would be more efficient for oligogenic traits and maybe less effective for polygenic traits in which GWAS only detected a few SNP-trait associations with limited PVE.

### Family size matters for the efficiency of genomic prediction

Model training using higher numbers of clones per family, is usually expected to capture more Mendelian segregation effects [43] and improves the PA and accuracy of genetic parameter estimates [44]. In this study, we observed that increasing the family size was important to improve the PA, both with respect to overall phenotypic prediction (Fig. 2) but also to within-family prediction (Fig. 4). This was especially obvious when the family size was small (less than 15 clones per family).

### Relationship between training and validation datasets highly important for genomic prediction

Forward-selection tree breeding usually entails the selection of a few elite candidates within a progeny test population (F1) and the candidates are in turn crossed to produce a new batch of progenies (F2). Thus, when using existing F1 as a training dataset to predict F2, the average relationship between F1 and F2 will be lower compared with relationships within the same F1 generation as was used for 10-fold random cross-validations in this study. We therefore also investigated the situation where training and validation datasets contained individuals from separate families, and we produced a 32-fold cross-validation scheme by removing phenotypic data for one family at a time. The relationships between the validation family and the training dataset were thus restricted to the level of half-sibs or weaker. Thus, the PA across families was considerably lower than for conventional random cross-validation (Table 4), which was a result similar to that in *Pinus taeda* L. [35]. However, it is notable that the seemingly oligogenic BB still offered an appreciable PA estimate (0.19) and this estimate was further improved if a marker preselection was performed for calculating the *G_a_*-matrix of the model (GBLUP-S, 0.25) or by fitting the most significant marker as a fixed effect in the model (GBLUP-F, 0.24).

### Marker density and LD affects predictive ability of phenotypic variation and within-family variation

Marker density (i.e. the number of markers) is usually considered as one of the most important factors affecting GP performance [45]. However, in forest tree breeding, the capture of the expected pedigree relationships (e.g. 0.25 among half-sib siblings), only requires a few thousand SNPs. Literature indicates that such a number of markers would also be enough to achieve an overall predictive ability similar to models utilizing all markers in the same batch of markers [13, 17, 46]. In such situations, a few thousand SNPs preselected based on GWAS or other prior information could instead improve the PA, especially for traits where several large-effect QTNs have been found [17, 19, 40]. In theory, coefficients of the pedigree relationship matrix describe additive genetic relationships between individuals at quantitative traits loci [47], but in reality, it is not obvious to what extent the genomic relationship matrix explains a genetic covariance matrix between individuals for QTLs, especially for a targeted trait.

In animal breeding, for example in cattle breeding, using imputed whole-genome markers seems to capture similar phenotypic variation as using ~60k SNP array due to strong LDs between makers and QTL within a few breeds and also a moderate genome size, ca. 3.1G [5]. However, conifers usually have a much large genome size, such as Norway spruce with 20G of the total genomic content [48]. And also, LD decay in the tree-breeding population is much faster than in the cattle-breeding population. In this study, the extent of LD (*r*^2^ ≥ 0.2) was observed to be 42.9kb when based on the 50k SNP array used in this population (Fig. S2). Based on such an extent of LD between markers and QTLs, the 50k SNP array only covered ca. 25% of genome size. This could be one of the reasons why the PA of GP did not reach the standard value of the square root of additive clone mean heritability (Table 2) and why within-family and across-family PA was low for most of the studied traits (Table 4 and Fig. 4). In order to capture more Mendelian segregations between QTLs and makers, we would then need more markers for a successful marker preselection for oligogenic traits and to increase the number of informative markers for polygenic traits in general. For BB, as an example, the model with 100 pre-selected markers based on GWAS may have captured LD between markers and QTLs, and indeed captured a considerable amount of within-family variation in agreement with a few studies on other tree species [17, 19, 49].

## Conclusion

Genomic selection with a marker preselection is considered an efficient approach in animal and tree breeding. For the budburst stage (BB), indicated as the oligogenic trait in this study, a preselection of approximately 100 SNPs based on the smallest *p*-values from GWAS,. showed the highest predictive ability (PA). But for the other polygenic traits, approximate 2000-4000 preselected SNPs, indicated by the smallest Akaike information criterion to offer the best model fit, still resulted in PA being similar to that of GP models using all markers. Analyses on both real-life and simulated data also showed that the inclusion of a large QTL SNP in the model as a fixed effect could improve PA and the accuracy of GP provided that the PVE of the QTL was ≥2.5%. Currently, most of the published marker resources designed for genomic selection in tree species, such as exome capture, genotyping-by-sequencing (GBS), and commonly designed SNP arrays did not consider or regardless of the marker density within strong LD regions. Therefore, within such a marker panel, many markers could be in strong LDs and not related to QTLs. So all markers used in the genomic model will not necessarily improve genomic prediction, and may even decrease the prediction power, such as our studied trait BB. Thus, we encourage performing a marker preselection step for genomic prediction, especially when those whole genome sequencing data or whole genome-imputed markers data may be available in the future. Meanwhile, the inclusion of a large QTL SNP in the model as a fixed effect is also strongly recommended in genomic selection.

## Materials and methods

### Plant materials

A Norway spruce breeding population using 32 control-pollinated families from 49 parents was established in 2007 at four different field sites. A total of 1430 unique clones were derived from the 32 families with an average of about 45 clones per family and about three ramets per genotype were originally planted in each site. The detailed descriptions of the four sites in this breeding population were presented in the published paper [21]. Generally, the field site series were established using a randomized incomplete block design with single-tree plots. Meanwhile, 98% of clones were replicated among the four sites. In this study, we selected all available clones from ten families (ca. 50 clones per family) and 20 clones from each of the remaining 22 families (a total of 904 clones) from a single site S1389 (Rössjöholm) for genotyping (Table S5).

### Phenotyping

Tree height was measured at field age six (HT6) at all four sites and at twelve years (HT12) at one of the sites. Diameter at breast height (DBH) was measured at field age twelve at all four sites. Budburst stage (BB) was scored at field age six at three sites based on eight categories [50]. Two further traits were measured only in a single site. Pilodyn penetration (PILO) as a proxy of wood density was measured by Pilodyn 6J Forest (PROCEQ, Zurich, Switzerland) at field age twelve. Frost damage (FD) was quantified after the site was exposed to a severe frost event at field age six. FD was scored as a categorical variable from zero (without frost damage) to three (the most severe damage). The detailed descriptions of traits measured in each of the four sites are shown in Table S6.

### Genotyping

Newly fresh needles were sampled from 904 clones in the spring of 2018. Total genomic DNA was extracted using the Qiagen plant DNA extraction protocol with DNA quantification performed using the Qubit® ds DNA Broad Range Assay Kit (Qiagen, Oregon, USA). Genotypic data were generated using the Norway spruce *Piab50K* SNP array chip [51]. Genotype calling of the 50k Axiom array was performed as the description in the paper [51]. Here, missing SNPs were imputed by Beagle v4.0 [52]. Due to several parents only involving a single controlled cross/family, the unique rare allele of the parent only contributes one allele to each progeny in the family. Thus, SNPs with a minor allele frequency (MAF) less than *M*/(2*n*) were filtered out for all models, where *M* is the harmonic number of clones per family and *n* is the population size in each GP model.

### Pedigree correction

Since a couple of clear discrepancies between the additive relationship and the genomic relationship matrices (***A*** and ***G_a_***, respectively) were detected for some individuals, we performed a pedigree correction for this population based on ***A*** and a heatmap of ***G_a_***. The number of parents increased from 49 based on the documented pedigree to 55 based on ***G_a_*** and the number of families also increased from the original 32 to 56 (Table S5). Finally, the number of clones per family after correction varied from 1 to 56.

### Spatial analysis

Spatial analysis based on a two-dimensional separable autoregressive (AR1) model was used to fit the row and column directions for phenotypic data from each site using ASReml v4.1 (Gilmour *et al*, 2015). Block effects were estimated simultaneously. Adjusted data, where all significant block and spatial effects were removed, were used for downstream analyses.

### Variance component and heritability estimates

Four univariate models were used to estimate variance components for each trait based on pedigree-based best linear unbiased prediction (PBLUP) and genomic-based best linear unbiased prediction (GBLUP) as following:

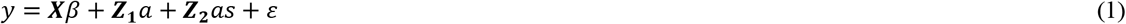

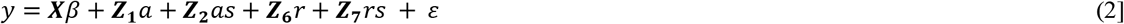

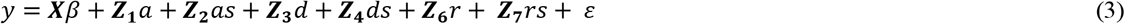

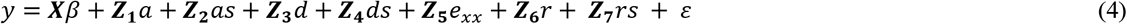

where ***y*** is the vector of adjusted phenotypic observations of a single trait; *β* is the vector of fixed effects, including a grand mean and site effects; *a* and *d* are the vectors of random additive and dominance effects, respectively; *as* and *ds* are the vectors of random additive-by-site and dominance-by-site effects, respectively; *e_xx_* is one of *e_aa_, e_ad_, and e_dd_*, which are the vectors of random additive-by-additive, additive-by-dominance, and dominance-by-dominance epistatic effects, respectively; *r* is the vector of residual genotypic effects, referring to an un-dissectable combination of dominance and epistatic effects in equation (2), epistatic effects in equation (3), epistatic effects excluding *e*_xx_ effects in equation (4); *rs* is the vector of residual genotypic-by-site effects; *ε* is the vector of random residual effects. ***X, Z*_1_, *Z*_2_, *Z*_3_, *Z*_4_, *Z*_5_, *Z*_6_**, and ***Z*_7_** are the incidence matrices for *β, a, as*, *d, ds, e_xx_*, *r*, and *rs*, respectively (detailed descriptions in Supplementary S1).

Pedigree-based BLUP (PBLUP) models based on equations (1) to (4) are called as PBLUP-A, PBLUP-AR, PBLUP-ADR, and PBLUP-ADR-xx, respectively. PBLUP-ADR-xx included three models of different epistatic effects called as PBLUP-ADR-aa, PBLUP-ADR-ad, and PBLUP-ADR-dd, respectively. Genomic-based BLUP models based on questions (1-4) could be called GBLUP-A, GBLUP-AR, GBLUP-ADR, and GBLUP-ADR-xx, respectively.

The pedigree-based additive (***A***) and dominance (***D***) relationship matrices were produced using the AGHmatix package [53]. The genomic-based additive (***G*_a_**) and dominance (***G*_d_**) relationship matrices were constructed based on imputed SNP data as described by [54] for ***G*_a_** and by [55] for ***G*_d_** using AGHmatrix package in R [53]. The relationship matrices due to the first-order epistatic interactions were computed using the Hadamard product (cell by cell multiplication, denoted #) and trace (*tr*) [56]. More detailed description of the matrices are shown in Supplementary methods S2 whereas the estimates of genetic variance parameters are shown in Supplementary methods S3

### Association mapping

To check the additive genetic architecture of the six traits in the breeding population, we also performed GWAS based on clone mean values across-site for all traits using the multi-locus BLINK model [57] conducted in GAPIT V3.0 R Software package [58]. Principal components were used to control population structure if the genomic inflation factor (i.e. lambda) is less than 0.95 or more than 1.05 [59]. The genome-wide significance of associations was determined at an experiment-wise false discovery rate 0.05 according to Benjamini & Hochberg (1995) [60]. The percentage of phenotypic variance explained (PVE) for each significant association was obtained from results using a mixed linear model (MLM) conducted in GAPIT V3.0 R software package [58].

### Linkage disequilibrium

Genome-wide analysis of linkage disequilibrium (LD) was conducted in the F1 full-sib progeny population. All SNPs were mapped into Norway spruce genome v2.0 (In preparation). Out of the 47,445 SNPs available from the *Piab50K* SNP array, 43,267 SNPs were successfully mapped and evenly distributed across the 12 chromosomes (Fig. 1a). The remainder 4178 SNPs could not be correctly mapped to any chromosome, therefore they were instead grouped together in an assumed “chromosome 13” (Table S2). LD values for pair-wise SNPs within each chromosome were calculated using VCFtools [61].

### Cross-validation test

Due to the negligible effects of dominance, dominance-by-site, and also first-order epistatic effects for all traits (Table S1), clone means calculated across-site were used as phenotype values to perform cross-validations. Ten sets of 10-fold cross-validations were performed. In summary, we employed a model as below:

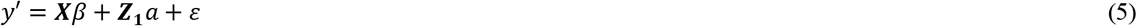

Where *y’* is the vector (904, 1) of the clonal means across-site, *β* is the vector of fixed effects (grand mean and an eventual single-locus SNP effect), *a* is the vector of additive effects and *ε* is the vector of random residual effects. ***X*** and ***Z*_1_** are the matrices related to the *β* and *a*. The random additive effects (*a*) were assumed to follow 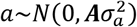 with 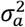 the additive variance in PBLUP-A whereas the matrix ***A*** will be replaced by ***G_a_*** in the GBLUP-A model.

The prediction ability (PA) was defined as the Pearson correlation between predicted breeding values (EBVs) and clonal means. Furthermore, we calculated the PA for within-family predictions as Pearson correlations between the EBVs deviation from EBV family mean and the clonal mean deviation from the family means used as benchmark validation values. An estimate of the selection accuracy was calculated by dividing PA with the square root of the pedigree-based narrow-sense clonal mean heritability (PA/*h_c_*).

### Testing the efficiency of genomic prediction

#### Marker density and maker preselection

To test the impact of the number of SNPs on the overall and within-family PA of GBLUP, we performed GPs using 14 subsets of SNPs (25, 50, 100, 200, 500, 1K, 2K, 4K, 8K, 10K, 20K, 30K, 40K, and all SNPs) and using two different types of sampling strategies: 1) randomly selected SNP subsets and 2) SNP subsets selected based on the smallest *p*-values shown in the GWAS for additive effects using BLINK method in the training population. We performed these steps in both the whole population with 10-fold cross-validation replicated 10 times (*n*=100) and in a subset population with 482 clones from the 10 largest families. The subset population was evaluated by inspecting their Aikake Information Criterion values (AIC) and the subset sample of markers showing the smallest AIC-values were selected for further investigations (henceforth called GBLUP-S).

#### Different statistical models

In most conifer tree species, including Norway spruce [40], growth traits, such as tree height and DBH, are commonly assumed as polygenic traits, but phenology traits, such as budburst stage may be considered as an oligogenic trait. Thus, we tested the efficacy for four different statistical models:

1. PBLUP-C: the traditional pedigree-based BLUP including pedigree correction.
2. GBLUP: a genome-based BLUP with a relationship matrix (***G_a_***) estimated from all markers.
3. GBLUP-S: a genome-based BLUP with a ***G_a_*** matrix estimated from a subset of preselected markers selected based on GWAS results generated for each training population separately. The number of preselected markers depended on the genetic architecture of traits.
4. GBLUP-F: a genome-based BLUP with a ***G_a_*** matrix estimated from all markers, except the marker with the greatest significance (smallest *p*-value) included as a fixed regression effect. This single marker was selected from GWAS results based on each training population.

In addition to the four models mentioned above, we also performed several extra strategies.

#### Relationship between training and validation sets

Compared with the previously random cross-validation strategy, another validation method entailed the prediction of genomic breeding values of all progenies within a specific full-sib family based on model training using all the other families using GBLUP. In all, there were 32 such across-family cross-validation repeats and the relationships between training and validation sets were thus consistently weak (unrelated or half-sib). The PA within the validated family therefore largely depends on whether the model could capture the Mendelian segregation effects.

#### Family size (i.e. number of clones per family)

To test if an increase in family size could improve the PA of PBLUP-C and GBLUP, we randomly selected five to 30 clones per family as a training set for the largest ten families with 48-56 clones per family (Fig. S3), using the remaining clones in the ten families as a validation set. This evaluation was performed both for regular PA and within-family PA and cross-validation was performed based on the corrected pedigree where the number of clones per family varied from 32 to 56 (Table S5).

### Simulations of large-effect SNPs and their inclusion in the genomic prediction model

To verify whether the inclusion of a major-effect locus as a fixed effect in the model would improve PA, we conducted finite-locus model simulations of a simplified breeding population undergoing one generation of directional selection. We used the software Metagene [62] to simulate a genomic architecture of 15,000 biallelic loci distributed along 12 chromosomes roughly like that of Norway spruce (see Supplementary methods S4 for details). The simulated breeding population comprised the crossing of 50 founders according to a single-pair-mating design producing 2000 offspring individuals in the F1-testing population. The heritability of the studied virtual trait was kept at 0.25 and an additional “major-gene locus” was introduced into simulations where the PVE of this locus was set throughout the range 0% (false-positive QTL), 0.25% (minor effect), 1.25%, 2.5% and 5% (major true QTL). GP-models (according to equation 1) were trained and predictions were cross-validated according to the methods described in previous subsections. One set of models where the presumed major-effect locus was included as a fixed regression effect (GBLUP-F) was compared to a corresponding set of models were the major-effect locus was not included (GBLUP).

## Acknowledgements

We greatly appreciated that Dr. Wu Yanfang help us with field samplings. The computations were performed on resources provided by the Swedish National Infrastructure for Computing (SNIC) at UPPMAX.

## Declarations

### Availability of data and materials

The phenotypic datasets supporting the conclusions of this article are available upon request to corresponding authors Z-QC and HXW. No voucher specimen has been deposited in this study. The genomic data generated and/or analysed during the current study are available in the zenodo repository, [https://doi.org/10.5281/zenodo.4781376.]

### Authors’ contributions

Z-QC, HRH, and HXW designed the study. Z-QC collected the needles in the field station. HRH and AK provided the long-term field phenotypic data. HRH produced the simulation data. Z-QC wrote the first draft of the manuscript whereas Z-QC, HRH, AK, and HXW made further revisions. HRH and HXW provided the funding for field data collection, genotyping, and data analysis.

### Ethics approval and consent to participate

The plant materials analyzed for this study come from common garden experiments (Plantation and clonal archives) that were established and maintained by the Forestry Research Institute of Sweden (Skogforsk) for breeding selections and research purposes. Two tree breeders, HRH and AK in Sweden were coauthors in this paper. They agreed to access the materials and collect the needles. Experimental research and field studies on plants including the collection of plant material are comply with relevant guidelines and regulation.

### Consent for publication

Not applicable

### Author details

^1^Umeå Plant Science Centre, Department Forest Genetics and Plant Physiology, Swedish University of Agricultural Sciences, SE-90183 Umeå, Sweden. ^2^Skogforsk, SE-91821 Sävar, Sweden. ^3^Skogforsk, Uppsala Science Park, SE-75183 Uppsala. ^4^CSIRO National Collection Research Australia, Black Mountain Laboratory, Canberra, ACT 2601, Australia.

## Legends of supplementary Figures

**Fig. S1.**
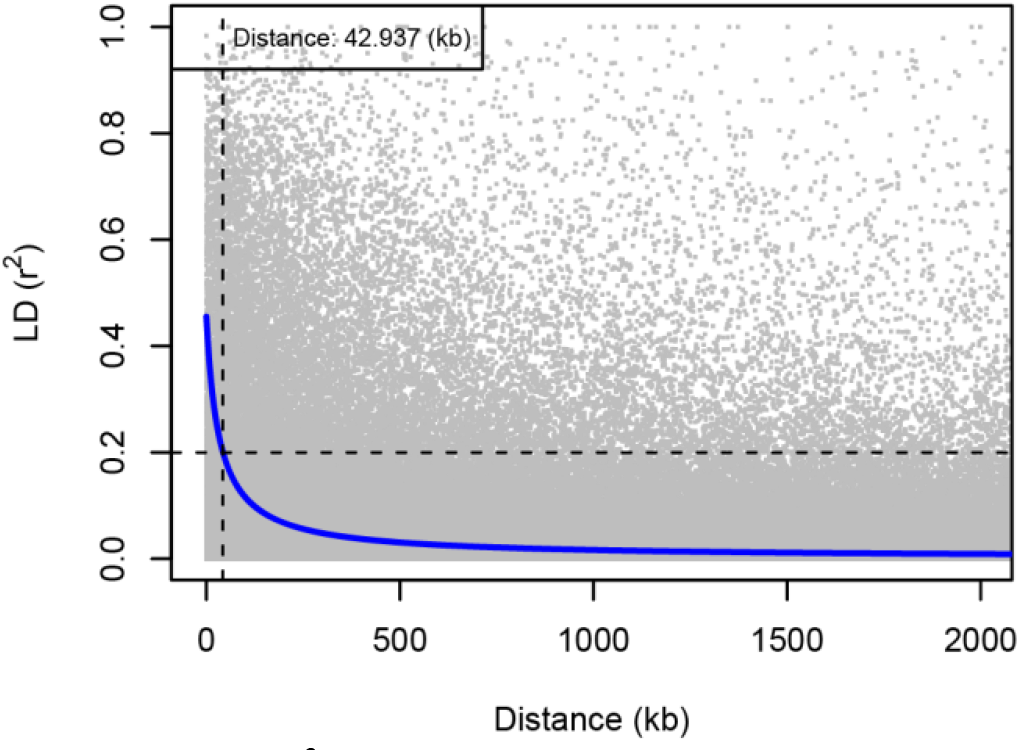
LD decay (*r*^2^) in a full-sib progeny population (F1-generation, n=904 from 49 parents based on the original pedigree) based on Norway spruce 50k SNP array. The Vertical dashed line represents LD decay distance when *r*^2^=0.2.

**Fig. S2.**
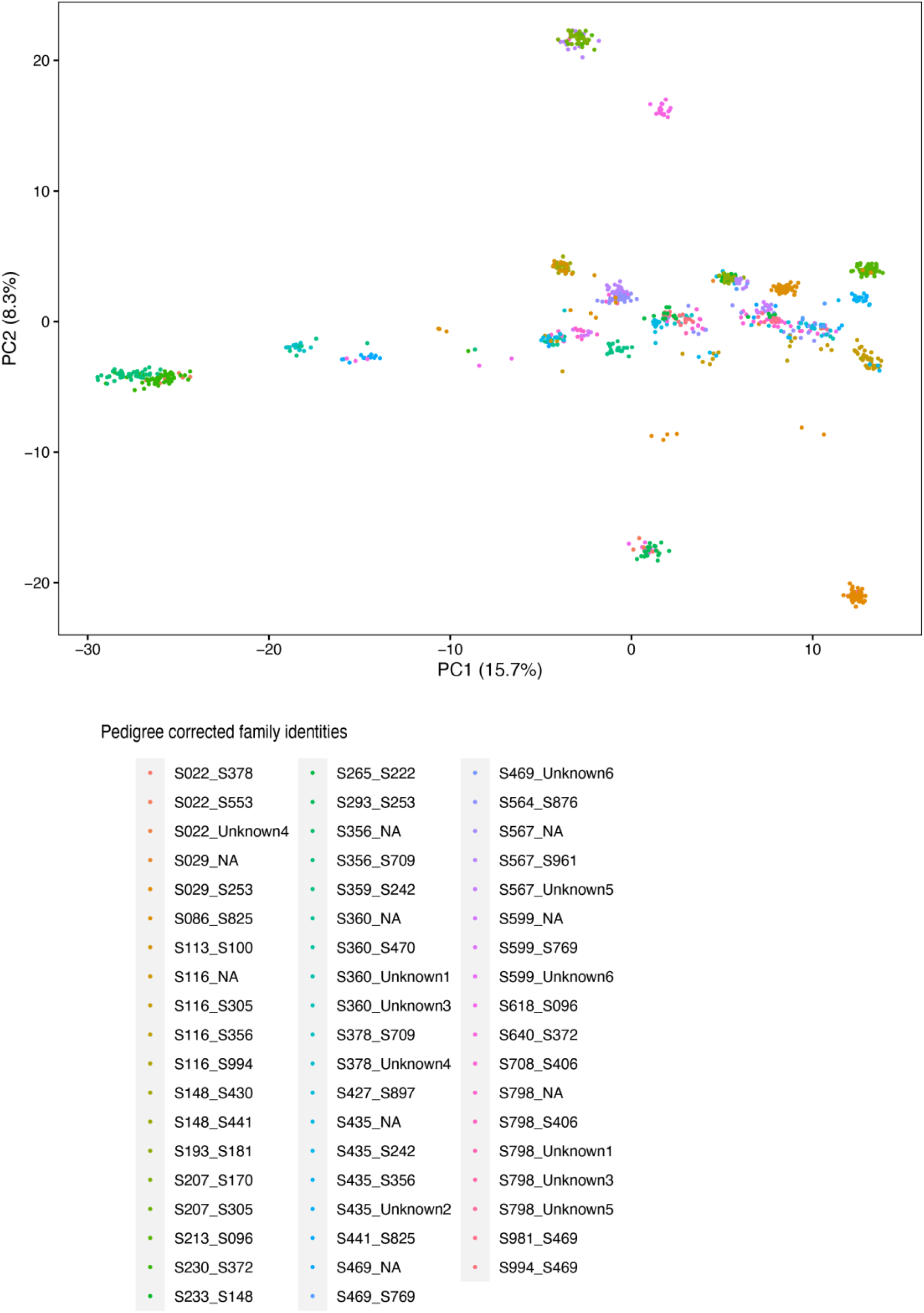
Population structure for the 904 clones coloured based on the family identities.

**Fig. S3.**
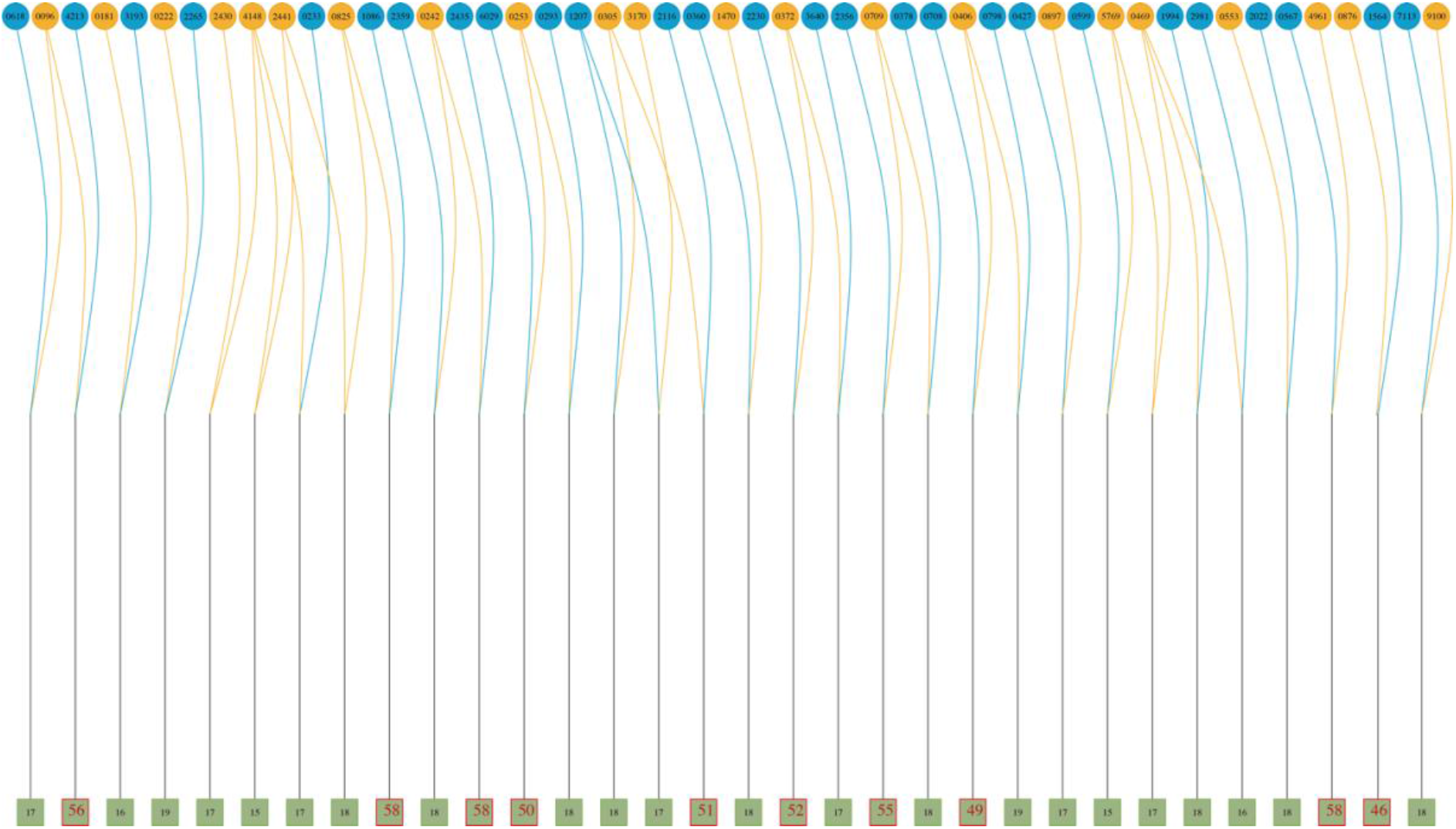
A mating design of 49 parents constructed into 32 full-sib families. Each circle in the top level represents a parent and the number is the identity of the parent. Each square at the bottom represents a full-sib family. The number within each square represents the number of clones per family genotyped. The families whose squares are framed in red colour, were selected to test the effect of family size on predictive ability. If the parents only cross once, then dark goldenrod means male and dark skyblue means female. Otherwise, the colour of the parents could be any of both colours.

