## Supplementary Methods 1-4 for "Preselection of QTL markers enhances accuracy of genomic selection in Norway spruce"

**Supplementary methods S1:**

Four univariate models were used to estimate variance components for each of three traits (HT6, DBH, and BB) based on pedigree-based best linear unbiased prediction (PBLUP) and genomic-based best linear unbiased prediction (GBLUP) as following:

$$y = \boldsymbol{X}\beta + \boldsymbol{Z_1}a + \boldsymbol{Z_2}as + \varepsilon \quad (1)$$

$$y = \boldsymbol{X}\beta + \boldsymbol{Z_1}a + \boldsymbol{Z_2}as + \boldsymbol{Z_6}r + \boldsymbol{Z_7}rs + \varepsilon \quad (2)$$

$$y = \boldsymbol{X}\beta + \boldsymbol{Z_1}a + \boldsymbol{Z_2}as + \boldsymbol{Z_3}d + \boldsymbol{Z_4}ds + \boldsymbol{Z_6}r + \boldsymbol{Z_7}rs + \varepsilon \quad (3)$$

$$y = \boldsymbol{X}\beta + \boldsymbol{Z_1}a + \boldsymbol{Z_2}as + \boldsymbol{Z_3}d + \boldsymbol{Z_4}ds + \boldsymbol{Z_5}e_{xx} + \boldsymbol{Z_6}r + \boldsymbol{Z_7}rs + \varepsilon \quad (4)$$

where $\boldsymbol{y}$ is the vector of adjusted phenotypic observations of a single trait; $\beta$ is the vector of fixed effects, including a grand mean and site effects; $a$ and $d$ are the vectors of random additive and dominance effects, respectively; $as$ and $ds$ are the vectors of random additive-by-site and dominance-by-site effects, respectively; $e_{xx}$ is one of $e_{aa}, e_{ad}, and\ e_{dd}$, which are the vectors of random additive-by-additive, additive-by-dominance, and dominance-by-dominance epistatic effects, respectively; $r$ is the vector of residual genotypic effects, referring to an un-dissectable combination of dominance and epistatic effects in equation (2), epistatic effects in equation (3), epistatic effects excluding $e_{xx}$ effects in equation (4); $rs$ is the vector of residual genotypic-by-site effects; $\varepsilon$ is the vector of random residual effects. $\boldsymbol{X}$, $\boldsymbol{Z_1}$, $\boldsymbol{Z_2}$, $\boldsymbol{Z_3}$, $\boldsymbol{Z_4}$, $\boldsymbol{Z_5}$, $\boldsymbol{Z_6}$, and $\boldsymbol{Z_7}$ are the incidence matrices for $\beta$, $a$, $as$, $d$, $ds$, $e_{xx}$, $r$, and $rs$, respectively. The random additive effects $(a)$ in equation (1–4) were assumed to follow $a \sim N(0, \boldsymbol{A}\sigma_a^2)$ with $\sigma_a^2$ the additive variance, where $\boldsymbol{A}$ is the pedigree-based additive relationship matrix in PBLUP (or replaced by a genomic-based additive relationship matrix $\boldsymbol{G_a}$ in GBLUP). The random dominance effects ($d$) in equations (1–4) were assumed to follow $d \sim N(0, \boldsymbol{D}\sigma_d^2)$ with $\sigma_d^2$ the dominance variance, where $\boldsymbol{D}$ is the pedigree-based dominance relationship matrix in PBLUP (or replaced by a genomic-based dominance relationship matrix $\boldsymbol{G_d}$ in GBLUP). The $as$ and $ds$ are the random additive-by-site and dominance-by-site interaction effects following $as \sim N(0,\ \sigma_{as}^2 \boldsymbol{I_s} \otimes \boldsymbol{A})$, and $ds \sim N(0, \sigma_{ds}^2 \boldsymbol{I_s} \otimes \boldsymbol{D})$ in the PBLUP model (or replaced by $\boldsymbol{G}_{\mathbf{a}}$ and $\boldsymbol{G}_{\mathrm{d}}$ in the GBLUP model). $\sigma_{as}^2\ and\ \sigma_{ds}^2$ are the additive-by-site and dominance-by-site variances, respectively. $e_{aa}, e_{ad}, and\ e_{dd}$ are the vectors of the random additive-by-additive, additive-by-dominance and dominance-by-dominance epistatic effects following $e_{aa} \sim N(0, \boldsymbol{P_{aa}}\sigma_{aa}^2)$, $e_{ad} \sim N(0, \boldsymbol{P_{ad}}\sigma_{ad}^2)$, and $e_{dd} \sim N(0, \boldsymbol{P_{dd}}\sigma_{dd}^2)$, respectively. $\boldsymbol{P_{aa}}$, $\boldsymbol{P_{ad}}$, and $\boldsymbol{P_{dd}}$ are the pedigree-based additive-by-additive, additive-by-dominance, and dominance-by-dominance epistatic relationship matrices in the PBLUP model ($\boldsymbol{G_{aa}}, \boldsymbol{G_{ad}}$, and $\boldsymbol{G_{dd}}$ in GBLUP model), respectively and $\sigma_{aa}^2$, $\sigma_{ad}^2$, and $\sigma_{dd}^2$ are their variance components. The $r$ and $rs$ are the vectors of residual genotypic effects and residual genotypic-by-site effects following $r \sim N(0, \boldsymbol{I}_{\boldsymbol{nc}}\sigma_r^2)$ and $rs \sim N(0, \boldsymbol{I}_{\boldsymbol{ncs}}\sigma_{rs}^2)$, respectively, where *nc* is the number of clones, *ncs* is the number of clones multiplied by the number of sites. The vector of residual *e* was assumed to follow

$$\varepsilon \sim \mathrm{N}(0, \begin{bmatrix} \boldsymbol{I}_{\mathrm{n1}}\,\sigma_{\mathrm{e1}}^2 & 0 & 0 & 0 \\ 0 & \boldsymbol{I}_{\mathrm{n2}}\,\sigma_{\mathrm{e2}}^2 & 0 & 0 \\ 0 & 0 & \boldsymbol{I}_{\mathrm{n3}}\,\sigma_{\mathrm{e3}}^2 & 0 \\ 0 & 0 & 0 & \boldsymbol{I}_{\mathrm{n4}}\,\sigma_{\mathrm{e4}}^2 \end{bmatrix}),$$

where $\sigma_{\mathrm{e1}}^2$, $\sigma_{\mathrm{e2}}^2$, $\sigma_{\mathrm{e3}}^2$, and $\sigma_{\mathrm{e4}}^2$ are the residual variances for sites 1, 2, 3, and 4, respectively; $\boldsymbol{I}_{\mathbf{n1}}$, $\boldsymbol{I}_{\mathbf{n2}}$, $\boldsymbol{I}_{\mathbf{n3}}$, and $\boldsymbol{I}_{\mathbf{n4}}$ are their identity matrices, and n1, n2, n3, and n4 are the number of individuals at each of the four sites, respectively.

For the other three traits (HT12, PILO, and FD), four similar models as equations (1–4) were used, except that the interaction terms with the site were excluded because those traits were only measured at a single site (Table 1).

**Supplementary methods S2:**

**Pedigree-based and genomic-based relationship matrix estimates**

The pedigree-based additive ($\boldsymbol{A}$) and dominance ($\boldsymbol{D}$) relationship matrices were constructed based on information from pedigrees. The diagonal elements (*i*) of the $\boldsymbol{A}$ were calculated as $A_{ii} = 1 + f_i = 1 + A_{gh}/2$, where $g$ and $h$ are the parents of the $i$th individual, while the off-diagonal element is the relationship between individuals $i$th and $j$th calculated as $A_{ij} = A_{ji} = (A_{jg} + A_{jh})/2$ [1]. In the $\boldsymbol{D}$ matrix, the diagonal elements were all one ($D_{ii} = 1$), while the off-diagonal elements between the individual $i$th and $j$th can be calculated as $D_{ij} = (A_{gk}A_{hl} + A_{gl}A_{hk})/4$, where $g$ and $h$ are the parents of the $i$th individual and $k$ and $l$ are the parents of the $j$th individual. $\boldsymbol{A}$ and $\boldsymbol{D}$ relationship matrices were produced using the AGHmatix package [2].

The genomic-based additive ($\boldsymbol{G}_{\mathbf{a}}$) and dominance ($\boldsymbol{G}_{\mathbf{d}}$) relationship matrices were constructed based on imputed SNP data as described by VanRaden (2008) [3] for $\boldsymbol{G}_{\mathbf{a}}$ and by Vitezica et al. (2013) [4] for $\boldsymbol{G}_{\mathbf{d}}$ using AGHmatrix package in R [2].

The relationship matrices due to the first-order epistatic interactions were computed using the Hadamard product (cell by cell multiplication, denoted #) and trace (*tr*) [5]. In the pedigree-based model, the additive-by-additive terms are calculated as $\boldsymbol{P}_{\mathbf{aa}} = \frac{\boldsymbol{A}\#\boldsymbol{A}}{tr(\boldsymbol{A}\#\boldsymbol{A})/n}$, additive-by-dominance terms as $\boldsymbol{P}_{\mathbf{ad}} = \frac{\boldsymbol{A}\#\boldsymbol{D}}{tr(\boldsymbol{A}\#\boldsymbol{D})/n}$, and dominance-by-dominance terms as $\boldsymbol{P}_{\mathbf{dd}} = \frac{\boldsymbol{D}\#\boldsymbol{D}}{tr(\boldsymbol{D}\#\boldsymbol{D})/n}$. The $n$ is the number of genotyped individuals. In the genomic-based relationship matrix models, additive-by-additive terms are calculated as $\boldsymbol{G_{aa}} = \frac{\boldsymbol{G_a}\#\boldsymbol{G_a}}{tr(\boldsymbol{G_a}\#\boldsymbol{G_{a)/n}}}$, additive-by-dominance terms as $\boldsymbol{G_{ad}} = \frac{\boldsymbol{G_a}\#\boldsymbol{G_d}}{tr(\boldsymbol{G_a}\#\boldsymbol{G_d})/n}$, and dominance-by-dominance terms as $\boldsymbol{G_{dd}} = \frac{\boldsymbol{G_d}\#\boldsymbol{G_d}}{tr(\boldsymbol{G_d}\#\boldsymbol{G_d})/n}$.

***Supplementary methods S3:***

**Proportion of variance component to phenotypic variance**

The narrow-sense heritability can be estimated as $h_i^2 = \sigma_a^2/\sigma_p^2$, using the models constructed above, $\sigma_p^2$ is the total phenotypic variance (e.g. in equation (3) and within a multi-site model, $\sigma_p^2 = \sigma_a^2 + \sigma_{as}^2 + \sigma_d^2 + \sigma_{ds}^2 + \sigma_r^2 + \sigma_{rs}^2 + \sigma_e^2$). Similarly, the dominance variance to the total phenotypic variance ratio was calculated as $d^2 = \sigma_d^2/\sigma_p^2$, the first-order epistatic variance to the total phenotypic variance ratio was calculated as $i^2 = \sigma_{xx}^2/\sigma_p^2$, the residual genotypic variance to the total phenotypic variance ratio was calculated as $r^2 = \sigma_r^2/\sigma_p^2$, and the broad-sense heritability was estimated as $H_i^2 = \sigma_g^2/\sigma_p^2$, where $\sigma_g^2 = \sigma_a^2 + \sigma_r^2$ is based on equation (2), $\sigma_g^2 = \sigma_a^2 + \sigma_d^2 + \sigma_r^2$ is based on equation (3), and $\sigma_g^2 = \sigma_a^2 + \sigma_d^2 + \sigma_{xx}^2 + \sigma_r^2$ is based on equation (4). Clone mean narrow-sense heritability was estimated as $h_c^2 = \sigma_a^2/\sigma_{\bar{p}}^2$, where $\sigma_{\bar{p}}^2 = \sigma_a^2 + \frac{\sigma_{as}^2}{s} + \sigma_d^2 + \frac{\sigma_{ds}^2}{s} + \sigma_r^2 + \frac{\sigma_{rs}^2}{s} + \frac{\sigma_e^2}{n}$ based on equation (3), *n* is the harmonic mean of the total number of ramets per clone and *s* is the harmonic mean number of sites in which each clone was represented by one or more ramets.

***Supplementary methods S4***

***Simulated genomic architecture***

To verify whether the inclusion of a major-effect locus as a fixed effect in the model would improve PA, we conducted finite-locus model simulations of a simplified Norway spruce breeding population undergoing one generation of directional selection. We would thereby demonstrate the value of identifying major-effect loci e.g. by GWAS in a context of genomic selection and breeding. We used the simulation software Metagene [6] to simulate one generation of genomic selection and breeding.

A simplified genomic architecture setup comprising 15,000 biallelic loci, uniformly distributed along 12 chromosomes, each with a genetic length of 250 cM, and thus similar to the architecture of Norway spruce [7], was used. Metagene regulated the genetic length of the genome by allowing completely random recombination between chromosomes at meiosis and thereafter randomly assigning 30 additional crossovers uniformly throughout the genome for each simulated meiosis event. Thus, a total genome length of 3,000 cM was achieved

(0.2 cM between each neighbouring pair of intrachromosomal loci). In order to emulate the highly skewed allele frequency distributions observed in many populations, allele frequencies for all loci ($p$) were drawn from a U-shaped probability distribution [8] while setting the effective population size parameter ($N_e$) equal to the number of simulated founders (100). For simplification, the founder haplotypes were sampled according to these specifications without the presence of any "historical" linkage disequilibrium (LD) thus implying that alleles would roughly adhere to Hardy-Weinberg equilibrium (HW-eq) and that the founder population would be devoid of any systematic substructures. Out of the 15,000 available loci, 500 loci were randomly designed to have minor additive effects ($\alpha_i$ for effective loci $I$ = 1…500) on a virtual quantitative trait of interest in accordance to the results of previous meta-analyses [9]. To emulate the usually observed distribution of a few loci with slightly larger effects and many loci with very small effects, the sizes of the 500 additive allelic effects were randomly drawn from a negative exponential distribution (Otto and Jones, 2000) with the rate parameter (λ) set at 1. To regulate the heritability of the virtual trait, the total additive genetic variance of the founder population was first estimated [10] as:

$$\sigma_a^2 = 2\sum_{i=1}^{500}\alpha_i^2 p_i(1-p_i) \qquad (6)$$

Since the founder population adheres to HW-eq and the founder population exhibited no population structure (no systematic LD), this simplified equation should be adequate for estimating $\sigma_a^2$. Thereafter the virtual phenotypic trait was constructed by adding random environmental noise effects from a normal distribution with a mean of 0 and a standard deviation of $\sqrt{3}\sigma_a$ to the additive genetic effects ($2\sum_{i=1}^{500}\alpha_i p_{ij}(1-p_{ij})$ of each individual *j*. Thus, a virtual quantitative trait showing a narrow-sense heritability ($h^2$) of ~0.25 at the founder level was achieved. It should be noted that the additive genetic variance in subsequent simulated generations was allowed to be influenced by any simulated events that could occur (selection effects, deviations from HW-eq and accumulation of LD) whereas the environmental noise variance ($\sigma_e^2$) was fixated at the level determined at the founder level ($G_0$).

***Simulation of a single-population breeding program***

The 100 founders were crossed 50 times according to a randomly allocated single-pair mating design (SPM) yielding 40 offspring per cross and in total 2000 individuals distributed within 50 full-sib families in a segregation population. For the study of conventional genomic selection, 1500 loci, randomly selected among the total 15,000 loci, were "genotyped" and were subsequently subjected to the same type of GBLUP-analysis and cross-validation procedure as described previously for the real-life data (see equation (5)). However, for the simulated data, phenotypic data was used directly for training and as a validation benchmark because the simulations did not include clonal replicates.

The evaluation of genomic prediction PA was done as previously described for the real-life data whereas prediction accuracy was calculated as the Pearson-correlation between predicted breeding values and true breeding values as the latter is reported by Metagene. Within-family prediction accuracy was also estimated as the Pearson-correlation between predicted breeding value deviations from the family mean of estimated breeding values (EBVs) and the true EBVs – true family EBV mean as the validation benchmark. For a conventional GP, 10 repeated simulations were performed completely resampling genomic architecture and founder population makeup according to previously reported specifications and for each such simulation, one set of 10-fold cross-validations was performed.

***Simulations of major-effect loci***

Apart from conventional GP, simulation scenarios were designed in which the presence of a "major-effect" locus was added. To produce such a locus, the previously designed set of genomic architectures was modified so that a major locus was chosen within the existing genomic architectures. The major- effect locus for each architecture was arbitrarily selected within chromosome 1 but the choice of the locus was nonetheless done according to a number of requirements. The major effect locus was required to: 1) not exhibit any causal effect in the original genomic architecture; 2) exhibit a close-to-intermediate allele frequency ($0.45 < p_{maj} < 0.55$); 3) be situated at least 10 positions (2 cM) away from any other effective locus in order to avoid the development of overly tight linkage with these; and 4) not be previously assigned as a "genotyped locus" in the original setup. When added, the number of effective loci for the modified scenario thus became 501 rather than 500 for the conventional scenario. The effect size of the major locus ($\alpha_{maj}$) was regulated in terms of the percentage of the genetic variance that it could explain (PGVE) thus being calculated as:

$$\alpha_{maj} = \sqrt{\frac{PGVE}{100} \cdot \frac{\sigma_a^2}{2 \cdot p_{maj}(1-p_{maj})}} \quad [7]$$

Then, the effect size of all other effective loci was adjusted (rescaled) as:

$$\alpha_{i,adj} = \alpha_i \sqrt{1 - \frac{PGVE}{100}}$$

so that overall $\sigma_a^2$ remained constant. Environmental variance and heritability were regulated as previously stated. Different PGVE-values for the major locus were tested in different scenarios (0%, 1%, 5%, 10% and 20%). Because the previously used heritability of 0.25 was retained, this translates to 0%, 0.25%, 1.25%, 2.5%, and 5%

in terms of PVE. It should be noted that PVE = 0% (no major locus effect) would be equivalent to the conventional genomic architecture and a PVE at 0.25% would only imply that the locus would mainly fall within the conventional range of the negative exponential sampling distribution (i.e. a minor effect locus). However, a major locus with a PVE at 1.25% would equal the very strongest QTL that sampling of 500 effects from a negative exponential distribution could produce and major loci with PVE-values at 2.5% and 5% would be considerably stronger than what could be thus sampled with any reasonable likelihood (i.e. a true major QTL).

Using all these genomic architecture setups, with the PVE of the major locus varying from 0% (fake) to 5% (true major QTL), the conventional genomic prediction procedure was first simulated. As the major locus was not assigned as being genotyped in the conventional cross-validation procedure, these analyses represent the situation where a major locus (of various strengths) is present in the genome but is missed in genotyping and thus not directly utilized by the GP models. But in addition to this, a second set of cross-validation analyses were made where the major QTL was indeed genotyped and directly used as a fixed regression term in model training and application. This would represent the situation where the QTL was indeed observed and subsequently utilized in GP model training. It should be noted that the context of this strategy varies according to the relative strength (PVE) of the major locus, ranging from the attempted use of a locus that is not truly causal or major (PVE 0-0.25%) to the utilization of a true major-effect locus (PGVE 2.5-5%). Consequently, no GWAS analysis was performed to determine which locus would be used for model training and cross-validation in contrast to the real-data analysis.
